# Structural and Functional Responses of Soil Microbial Communities to Biodegradable Plastic Film Mulching in Two Agroecosystems

**DOI:** 10.1101/650317

**Authors:** Sreejata Bandopadhyay, Henry Y. Sintim, Jennifer M. DeBruyn

**Author notes:** Corresponding author Jennifer DeBruyn, Department of Biosystems Engineering and Soil Science, University of Tennessee, 2506 E.J. Chapman Drive, Knoxville, TN 37996, USA.

## Abstract

Polyethylene (PE) plastic mulch films are used globally in crop production but incur considerable disposal and environmental pollution issues. Biodegradable plastic mulch films (BDMs), an alternative to PE-based films, are designed to be tilled into the soil where they are expected to be mineralized to carbon dioxide, water and microbial biomass. However inadequate research regarding the impacts of repeated incorporation of BDMs on soil microbial communities has partly contributed to limited adoption of BDMs. In this study, we evaluated the effects of BDM incorporation on soil microbial community structure and function over two years in two geographical locations: Knoxville, TN, and in Mount Vernon, WA. Treatments included four plastic BDMs, a completely biodegradable cellulose mulch, a non-biodegradable PE mulch and a no mulch plot. Bacterial community structure determined using 16S rRNA amplicon sequencing revealed significant differences by location and season. Differences in bacterial communities by mulch treatment were not significant for any season in either location, except for Fall 2015 in WA where differences were observed between BDMs and no-mulch plots. Extracellular enzyme rate assays were used to characterize communities functionally, revealing significant differences by location and sampling season in both TN and WA but minimal differences between BDMs and PE treatments. Limited effects of BDM incorporation on soil bacterial community structure and soil enzyme activities when compared to PE suggest that BDMs have comparable influences on soil microbial communities, and therefore could be considered an alternative to PE.

**Importance:** Plastic film mulches increase crop yields and improve fruit quality. Most plastic mulches are made of polyethylene (PE), which is poorly degradable, resulting in undesirable end-of-life outcomes. Biodegradable mulches (BDMs) may be a sustainable alternative to PE. BDMs are made of polymers which can be degraded by soil microbial enzymes, and are meant to be tilled into soil after use. However, uncertainty about impacts of tilled-in BDMs on soil health has restricted adoption of BDMs. Our previous research showed BDMs did not have a major effect on a wide range of soil quality indicators (Sintim et al. 2019); here we focus on soil microbial communities, showing that BDMs do not have detectable effects on soil microbial communities and their functions, at least over the short term. This informs growers and regulators about use of BDMs in crop production, paving a way for an agricultural practice that reduces environmental plastic pollution.

## 1. Introduction

Plastic mulch films are widely used in crop production systems to improve soil microclimate and surpress weeds, translating into increased crop yields and/or improved fruit quality. Some of the agronomic benefits of using plastic mulch films include reduction of weed pressure (1), conservation of soil moisture (2, 3), and moderation of soil temperature, among others. Low density polyethylene (PE) mulch has traditionally been favored by growers due to its many attractive properties such as easy processability, high durability, flexibility etc. (4, 5). However, PE does not readily biodegrade, and thus must be disposed at the end of the growing season, contributing to our global plastic waste problem (6, 7). Even when removed from a field, fragments of film are left behind in the soil, which can affect soil function and soil biota (8–13) or leach out into water systems and pollute aquatic ecosystems (14–19). As these plastics break down in soil, they form microplastics (20), contributing to terrestrial microplastic pollution (13, 20).

Plastic mulch use is expected to increase to meet increasing global food demands; therefore, it is imperative to find alternatives that will reduce the environmental footprint. Biodegradable mulch films (BDMs) are a potential alternative: BDMs are made of polymers that can be degraded by microbial action (21–24). In the field, BDMs perform like other plastic films by altering the soil microclimate and improving crop yields (25). However, unlike PE plastics, which require removal and disposal, BDMs are designed to be tilled into the soil where resident soil microbes are expected to degrade them over time. Under ideal circumstances, they should eventually be mineralized into carbon dioxide and water.

Despite being a promising sustainable alternative, adoption of BDMs has been limited (26). Currently available BDMs are not certified for use in organic crop production in North America as they are not 100% bio-based (27). Regulators are hesitant to allow these products until there is convincing evidence that they are safe for soil ecosystems. Thus, evaluating the impacts of incorporation of BDMs into soil on soil health is a critical part of adoption and policy development surrounding BDMs (28).

BDMs can impact soil health in two ways: indirectly, in a manner similar to PE films, by acting as a surface barrier to soil and modifying the soil microclimate, and directly, by addition of physical fragments and mulch carbon into soil after tillage (5). The body of research on the impacts of polyethylene films on soil microbial communities and functions can help us predict the indirect effect of BDMs on soil health. However, research on the direct effects of BDMs on soil microbial community structure and function remains poorly answered due to a dearth of research that directly compares BDMs and PE in the same study. Unless there is a direct comparison of BDMs and PE, it is difficult to tease apart whether the observed changes are above and beyond what you would expect from the application of PE mulch to the soil surface (5). These answers are critical if widespread use of BDMs is to be advocated. Previous studies have analyzed impacts of BDMs on soil microbial communities using PLFA profiling (29) and pyrosequencing (30) methods.

However, these studies did not use PE as a negative control so effects of BDM tilling on soil microbial community structure and function remain uncertain.

In this study, we evaluated the impacts of BDM use on soil microbial communities by directly comparing to PE mulch in a two-year vegetable crop field trial in two diverse climates (Knoxville, TN, in the southeastern USA and Mount Vernon, WA, in the northwestern USA). During this field trial, measurement of a suite of soil physiochemical properties and calculation of soil health indices revealed that the overall effect of mulching on soil health was minimal and that BDMs performed comparably to PE (31). The study by Sintim et al. (31) showed that the effect of location, time and their interactions were greater compared to the effects of mulch treatments. To build on this finding, we focused on biological soil health, evaluating the impacts of BDMs on 1) soil microbial community structure, characterized using 16S rRNA gene amplicon sequencing 2) soil microbial abundances, estimated using qPCR and 3) soil microbial community function, estimated by a suite of soil extracellular enzyme rates over the two-year field trial experiment. We tested the hypotheses that plastic mulches would significantly alter soil microbial community structure and function, but that there would be no significant differences between PE and BDM mulches.

## 2. Materials and methods

### 2.1 Plastic Mulch Films

Three commercially available biodegradable mulch films (BioAgri®, Naturecycle, Organix A.G. Film™,) and one experimental film comprised of a blend of polylactic acid (PLA) and polyhydroxyalkanoates (PHA) were tested alongside a polyethylene (PE) mulch (negative control), and cellulose mulch (WeedGuard Plus®, positive control). Physicochemical properties of mulches are reported in Table 1.

**Table 1.**
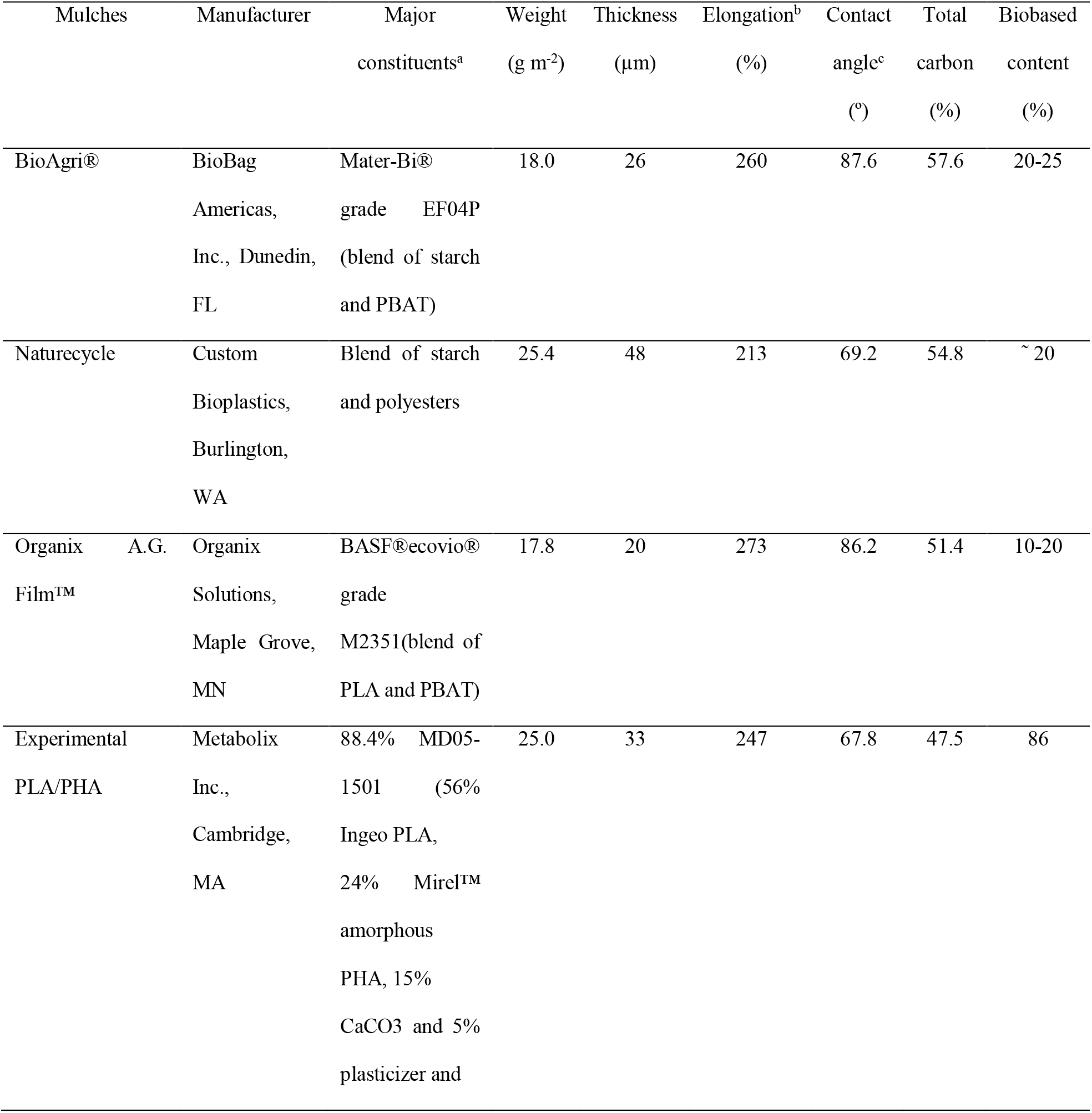
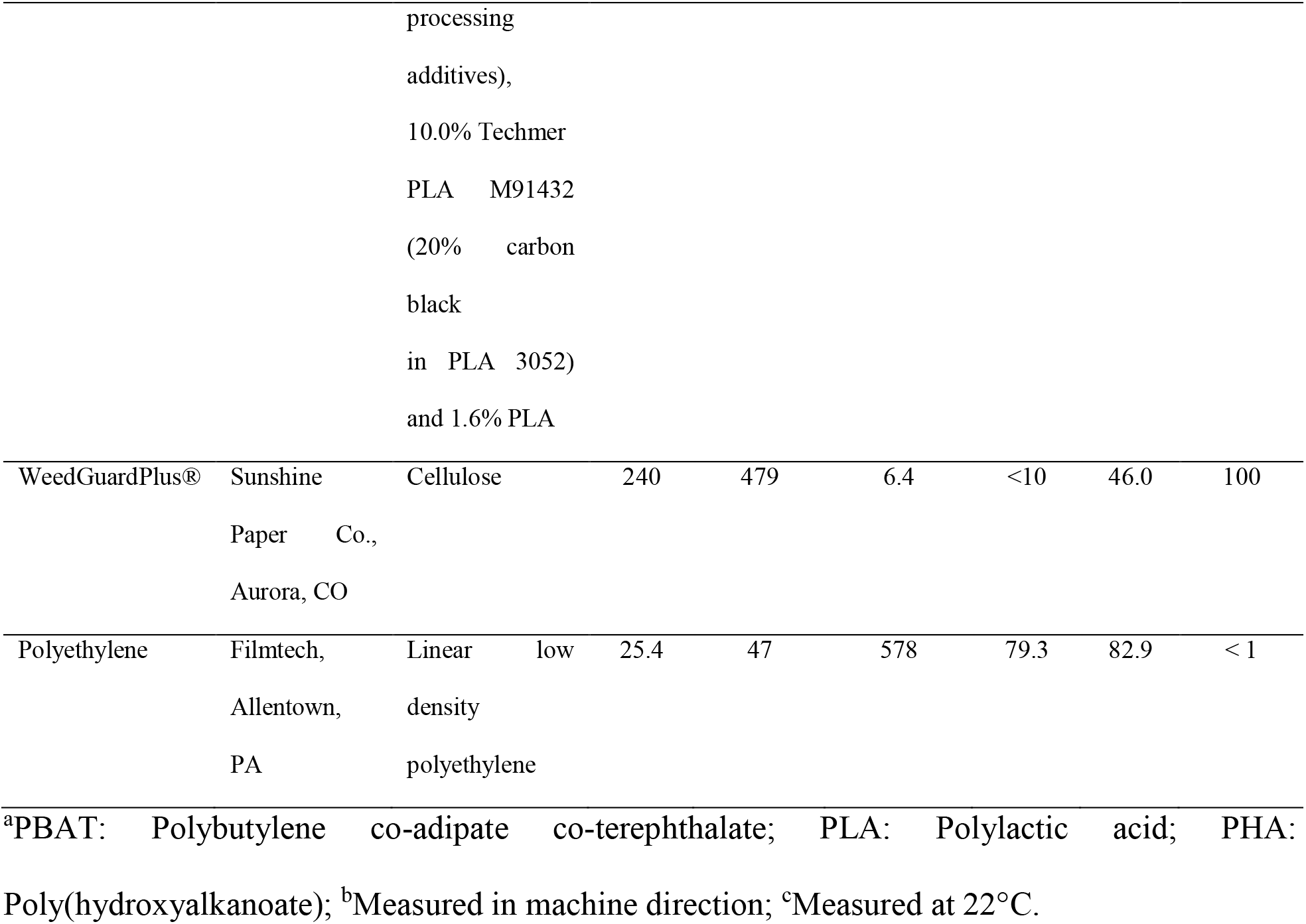
Manufacturers, major constituents, and physicochemical properties of the mulches used in the study. Bio-based content data was provided by the manufacturers. Data reported from Hayes et al. (51).

### 2.2 Field trial description

The mulches were tested in the field over two years (2015 to 2016) under pie pumpkin (*Cucurbita pepo*) as a test crop, with full experimental details described in Sintim et al. (31) and S. Ghimire et al. (32). Field experimental stations were set up in two locations: East Tennessee Research and Education Center (ETREC), University of Tennessee, Knoxville, TN and the Northwestern Washington Research & Extension Center (NWREC), Washington State University, Mount Vernon, WA. The soil at Knoxville is a sandy loam (59.9% sand, 23.5% silt, and 16.6% clay), classified as a fine kaolinitic thermic Typic Paleudults. The soil at Mount Vernon is a silt loam (14.2% sand, 69.8% silt, and 16% clay), classified as a fine-silty mixed nonacid mesic Typic Fluvaquents. Henceforth in the paper, Knoxville will be referred to as TN and Mount Vernon will be referred to as WA.

Each field site was arranged as a randomized complete block design with four replications of seven main plot treatments (six mulch treatments and one no mulch control). Before mulch application began in TN and WA, the plots were under winter wheat (*Triticum aestivum*) cover crop in TN and clover (*Trifolium spp.)* at WA. Mulches were machine-laid on raised beds. Pumpkins (*Cucurbita pepo*) were grown during the growing season. The PE mulch was removed after pumpkin harvest, while the BDMs were tilled into the soil with a rototiller.

Soil water content and temperature were monitored as described in Sintim et al. (31). Briefly, sensors (5TM, Decagon Devices Inc., Pullman, WA) installed in the center of each mulch treatment at 10-cm and 20-cm soil depths for one field block were connected to data loggers (EM50G, Decagon Devices Inc., Pullman, WA) that recorded the soil water and temperature data hourly. Soil water content and temperature data is reported in Sintim et al. (31). Air temperature, precipitation, relative humidity, wind, and solar radiation were collected from a meteorological station located at the field site at TN (Decagon Devices Inc. Weather Station, Pullman, WA), and about 100 m away from the field site at WA (WSU AgWeatherNet Station, Mount Vernon, WA). Weather data for the two locations for 2015-2017 are reported in Table S1.

Soil physical, chemical, and biological properties were assessed over the two-year study for this site, in order to assess changes in soil health. Detailed protocols for these measurements and raw data is provided in Sintim et al. (31).

### 2.3 Soil sampling

Soil samples were collected from each of the 28 plots (7 treatments, replicated 4 times) at both locations in the Spring (May) and Fall (September) of 2015 and 2016. Soil was collected from the top 10 cm, using a 2 cm diameter stainless steel auger. Thirty 10-cm soil cores were taken and composited for each of the plots. All sampling equipment was cleaned with 70% ethanol before and in between plots to limit cross contamination. Roots and pebbles were removed by hand, and soils homogenized and stored in plastic bags for transport back to the lab. Soils were stored at −80 °C until DNA extraction and extracellular enzyme assays.

### 2.4 Soil DNA extraction and quantification

Extraction of DNA from soil samples was completed using the MoBio™ PowerLyzer™ Power Soil DNA isolation kit (now branded under Qiagen™) with inhibitor removal technology, as per manufacturer’s instructions. 0.25 grams of soil were used for the extractions, and the DNA obtained after the final elution step was stored at −20 ⁰C until further analyses.

Quantification of the DNA extracted from soil was completed using the Quant-It™ PicoGreen™ dsDNA Quantification Kit (ThermoFisher Scientific) per manufacturer’s instructions. Standard curves generated had R squared values of 1. Mean DNA concentration of the soil samples was 13 ng µl^-1^ DNA.

### 2.5 Quantitative PCR for bacterial and fungal abundances

As a proxy for bacterial and fungal abundances, 16S rRNA (bacteria) and ITS (fungi) gene copy abundances were quantified from soil DNA samples using Femto™ Bacterial DNA quantification kit (Zymo Research) and Femto™ Fungal DNA quantification kit (Zymo Research) following the manufacturer’s protocol. DNA extracts were diluted 1:10 prior to quantification and 1 µl of the diluted samples was used for each qPCR reaction. All samples were analyzed in triplicate. No template negative controls were included in each run. Bacterial and fungal DNA standards were provided in the kit and the ng DNA standard per well was converted to copy numbers which were used for final calculations. qPCR reactions were performed in a CFX Connect Real-Time PCR Detection System (BioRad). qPCR efficiencies averaged around 85% and 90% for bacterial and fungal assays, respectively. Standard curves had R squared values ranging from 0.98 to 1.

### 2.6 DNA amplification and sequencing

16S rRNA amplicon sequencing of DNA extracts was conducted by the Genomic Services Laboratory (GSL) at Hudson Alpha, Huntsville, AL, following their standard operating procedures. Extracted DNA samples were shipped frozen in 96 well plates. The V4 region of the 16S rRNA gene was amplified using primers 515F (GTGCCAAGCAGCCGCGGTAA) and 806R (GGACTACHVGGGTWTCTAAT) (33). The first PCR was run with V4 amplicon primers, Kapa HiFi master mix, and 20 cycles of PCR. All aliquots and dilutions of the samples were completed using the Biomek liquid handler. PCR products were purified and were stored at −20⁰C until further processing was completed. The PCR indexing was later completed for the 16S (V4) amplicon batch. Products were indexed using GSL3.7/PE1 primers, Kapa HiFi master mix, and 12 cycles of PCR. Products were purified using magnetic beads using the Biomek liquid handler. Final libraries were quantified using Pico Green. V4 amplicon size obtained was 425 bp for the soil samples. The amplified 16S rRNA genes were sequenced using 250 paired-end reads on an Illumina MiSeq platform. Sequence reads are deposited in the NCBI sequence read archive (Accession XXXXXX).

Raw sequence data was processed using mothur v.1.39.5 following the MiSeq SOP (34) (Supplemental File 1). Before aligning to the reference database (SILVA release 102), unique sequences were identified, and a count table generated. After alignment to SILVA database, sequences were filtered to remove overhangs at both ends, and sequences de-noised by pre-clustering sequences with up to two nucleotide differences. Chimeras were removed using the VSEARCH algorithm. All sequences including 18S rRNA gene fragments and 16S rRNA from Archaea, chloroplasts, and mitochondria were classified using the Bayesian classifier (35) against the mothur-formatted version of the RDP PDS training set (v.9) with a bootstrap value of > 80% (35). Following this step, untargeted (i.e. non-bacterial) sequences classified as *Eukaryota* and *Arachaeota* were removed. Sequences were finally binned into phylotypes according to their taxonomic classification at the genus level. A consensus taxonomy for each OTU was generated by comparison to the RDP training set. The resulting OTU count table and taxonomy assignments were imported into R (v. 3.4.0) (36) for further downstream statistical analyses. Mothur code, R code and associated input files are available at: https://github.com/jdebruyn/BDM-Microbiology.

### 2.7 Extracellular enzyme assays

Fluorescence microplate enzyme assays were conducted using fluorescently labelled substrates to assess enzyme activities in soil (37). Seven enzymes were assayed using their respective fluorescent substrates and standards (Table 2).

**Table 2.**
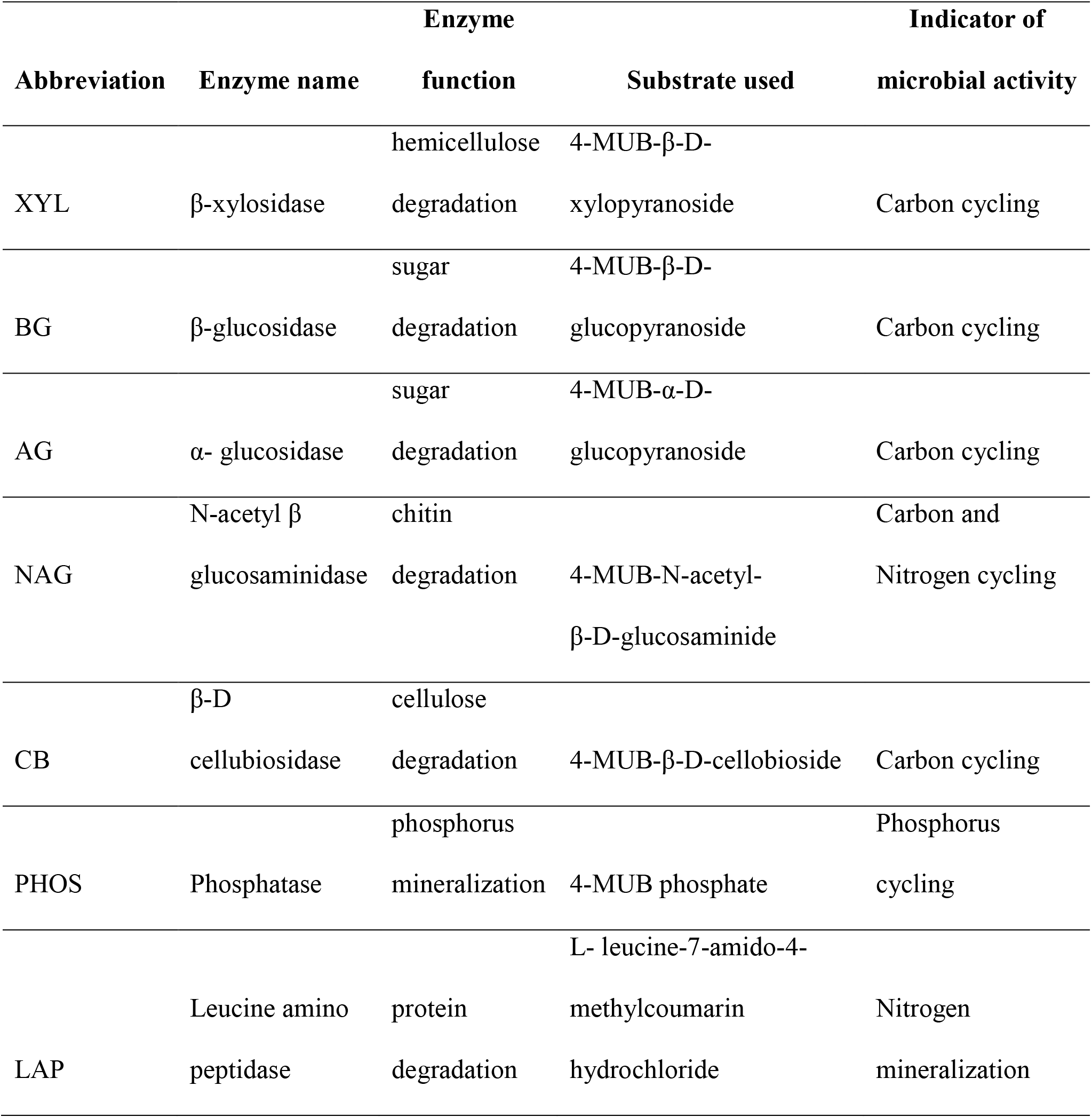
The seven extracellular soil enzymes assayed for soils collected from Spring 2015-Spring 2017; their respective enzyme functions, substrates used for assays, and the role of each enzyme in biogeochemical cycling. Standards used were MUB (4-methylumbelliferone) and MUC (7-amino-4-methylcoumarin).

| Abbreviation | Enzyme name | Enzyme |  | Indicator of microbial activity |
| --- | --- | --- | --- | --- |
|  |  | function | Substrate used |  |
| XYL | β-xylosidase | hemicellulose | 4-MUB-β-D- | Carbon cycling |
|  |  | degradation | xylopyranoside |  |
| BG | β-glucosidase | sugar | 4-MUB-β-D- | Carbon cycling |
|  |  | degradation | glucopyranoside |  |
| AG | α- glucosidase | sugar | 4-MUB-α-D- | Carbon cycling |
|  |  | degradation | glucopyranoside |  |
| NAG | N-acetyl $\beta$ | chitin | | Carbon and |
| | glucosaminidase | degradation | 4-MUB-N-acetyl-β-D-glucosaminide | Nitrogen cycling |
| CB | β-D | cellulose | | Carbon cycling |
| | cellubiosidase | degradation | 4-MUB-β-D-cellobioside | |
| PHOS | Phosphatase | phosphorus |  | Phosphorus |
|  |  | mineralization | 4-MUB phosphate | cycling |
| LAP | Leucine amino<br>peptidase | protein | L- leucine-7-amido-4-methylcoumarin | Nitrogen |
|  |  | degradation | hydrochloride | mineralization |

Soil slurries were prepared in a sodium acetate trihydrate buffer whose pH was matched closely with the soil pH. 800 µl of soil slurry was pipetted into deep well 96 well plates. Separate plates were prepared for MUB and MUC standard curves for each sample. 200 µl of appropriate standards and substrates were added to the soil slurries. The plates were sealed and inverted to mix the contents. Incubation was done for 3 hours at room temperature, after which the substrate and standard plates were centrifuged at 1500 rpm (∼327 x g) for 3 min. The supernatants were pipetted into black 96 well plates and fluorescence measured at 365 nm excitation wavelength and 450 nm emission wavelength in a BioTek® Synergy plate reader.

### 2.8 Statistical analyses

Beta diversity was computed using Bray-Curtis distances of microbial community composition using the vegan package (v 2.4-3) in R version 3.4.0 (36) based on OTU tables, and were then visualized using non-metric multidimensional scaling (NMDS) using phyloseq package v.1.21.0 in R (38). To determine whether significant differences existed in bacterial community composition between bacterial communities across different locations, seasons, and mulch treatments, a permutational multivariate analysis of variance (PERMANOVA) was performed using the ADONIS function implemented in R, based on the Bray-Curtis dissimilarity matrix. All libraries were scaled to even depth (minimum sample read count, i.e. smallest library size, of 34,266) before analysis was performed. Similarity percentage analyses (SIMPER) was completed in R to reveal the most influential OTUs driving differences between soil bacterial communities in different locations, and across different seasons. Canonical analysis of principal coordinates (CAP) was done to relate environmental variables reported in Sintim et al. (31) to changes in bacterial community composition. The ordination axes were constrained to linear combinations of environmental variables, then the environmental scores were plotted onto the ordination. A PERMANOVA was performed on the CAP axes. These analyses were completed in R following the online tutorial by Berry M (39).

Alpha diversity was computed by subsampling the libraries to the minimum number of reads (34,336). This was done with replacement to estimate species abundance of the real population by normalizing sampling effort. The subsampling was repeated 100 times and the diversity estimates from each trial were averaged. The estimate_richness function was used in R phyloseq package to calculate observed richness and inverse Simpson indices (for diversity). A mixed model analysis of variance was completed using the generalized linear mixed model (GLIMMIX) procedure in SAS V. 9.3 to assess changes in richness and inverse Simpson over time. The fixed effects were location (TN and WA), mulch treatments (7 treatments total) and date/season of soil sampling (4 time points), while random effect was block (total 3 blocks to serve as replicates). Repeated measures were incorporated in the model as sampling was done over time, twice a year in Spring and Fall seasons in 2015 and 2016. The model was a completely randomized design (CRD) split-split-plot with repeated measures in the sub-sub plot. Normality of data was checked using Shapiro-Wilk test (W > 0.9) and equal variance using Levene’s test (α = 0.05). All data were normal and hence no transformations were performed. Raw experimental values and standard errors are reported in the figures.

To visualize differences in the functional profile of the communities; i.e. all seven enzyme rates), NMDS ordination of Bray-Curtis similarities was done in Primer 7 v. 7.0.13 (PRIMER-E). A mixed model analysis of variance with repeated measures was completed using the generalized linear mixed model (GLIMMIX) procedure in SAS V. 9.3 to assess changes in enzyme activities over time. Fixed and random effects were same as specified above. However, location as a class was not included in this model as PERMANOVA results from PRIMER-E were used to report differences between locations. Boxplots for equal variance and outliers, reported in SAS, were used to remove outliers in the dataset. Normality was checked using Shapiro-Wilk test (W > 0.9) and probability plots for residuals, and equal variance using Levene’s test (α = 0.05). Data were log transformed as necessary when these conditions were not met. Raw experimental values and standard errors are reported in the figures. All graphics were plotted using R. v. 3.4.0. Type III tests of fixed effects and interaction effects are reported.

To assess for potential enrichment of bacteria and fungi, a paired t-test was conducted using initial and final 16S and ITS gene copy abundances (determined by qPCR) from Spring 2015 and Fall 2016 to see if there was a significant change. Initial 16S and ITS gene copy abundances from Spring 2015 were also subtracted from final abundances in Fall 2016 to get change in abundance over time. To determine if the enrichment or depletion of bacterial and fungal abundances was significantly different between treatments, a mixed model analysis of variance in SAS v. 9.3 using the GLIMMIX procedure was conducted on the differences. Significance level of all analyses were assessed at α = 0.05. All data were checked for normality using Shapiro-Wilk test (W > 0.9).

## 3. Results

### 3.1 Environmental and soil physicochemical data

Environmental data collected during the experiment is reported in Sintim et al. (31) and in Table S1. The mean daily air temperature in Knoxville, TN during experimental years of 2015 to 2016 was about 4 °C higher than in Mount Vernon, WA (Table S1). The total annual precipitation during the experimental years was higher in Knoxville, TN than in Mount Vernon, WA.

Soil temperature, moisture and physicochemical properties were measured and reported previously by Sintim et al. (31). In summary, significantly increased soil temperature was observed in the early growing seasons in the plastic mulch plots compared to the cellulose and no-mulch plots. On average, the monthly soil temperature was greater in TN than in WA. The soil water content varied more among the mulch treatments, with PE mulch having the highest soil water content for the greatest period of time. Mulched plots generally had higher water content than the no mulch plots. The soil health analysis revealed some effects of mulching on certain properties (namely aggregate stability, infiltration, soil pH, electrical conductivity, nitrate, and exchangeable potassium), but these were not consistent among BDMs, nor across sampling times and locations.

### 3.2 Soil bacterial community diversity and structure

The NMDS ordination revealed a clear difference in community structure between TN and WA when combining data from all four sampling seasons (Spring 2015 to Fall 2016) (Fig 1a). Permutational ANOVA (PERMANOVA) tests confirmed significant differences between TN and WA soil microbial communities (Table 3, Table S2). The mean relative abundances of the most abundant classes of bacteria are shown in Fig 1b. Similarity percentage tests (SIMPER) revealed the most influential OTUs contributing to the variation seen between location (Fig 1b). The most influential OTUs belonged to several classes of microbes such as *Acidobacteria_Gp7, Acidobacteria_Gp16, Acidobacteria_Gp4, Planctomycetacia* and *Spartobacteria.* CAP analysis revealed that the differences in soil communities between TN and WA were most related to pH, soil moisture and organic matter content: the communities in TN were related to increased pH, whereas moisture and organic matter were positively related to communities in WA (Fig S1).

**Fig 1.**
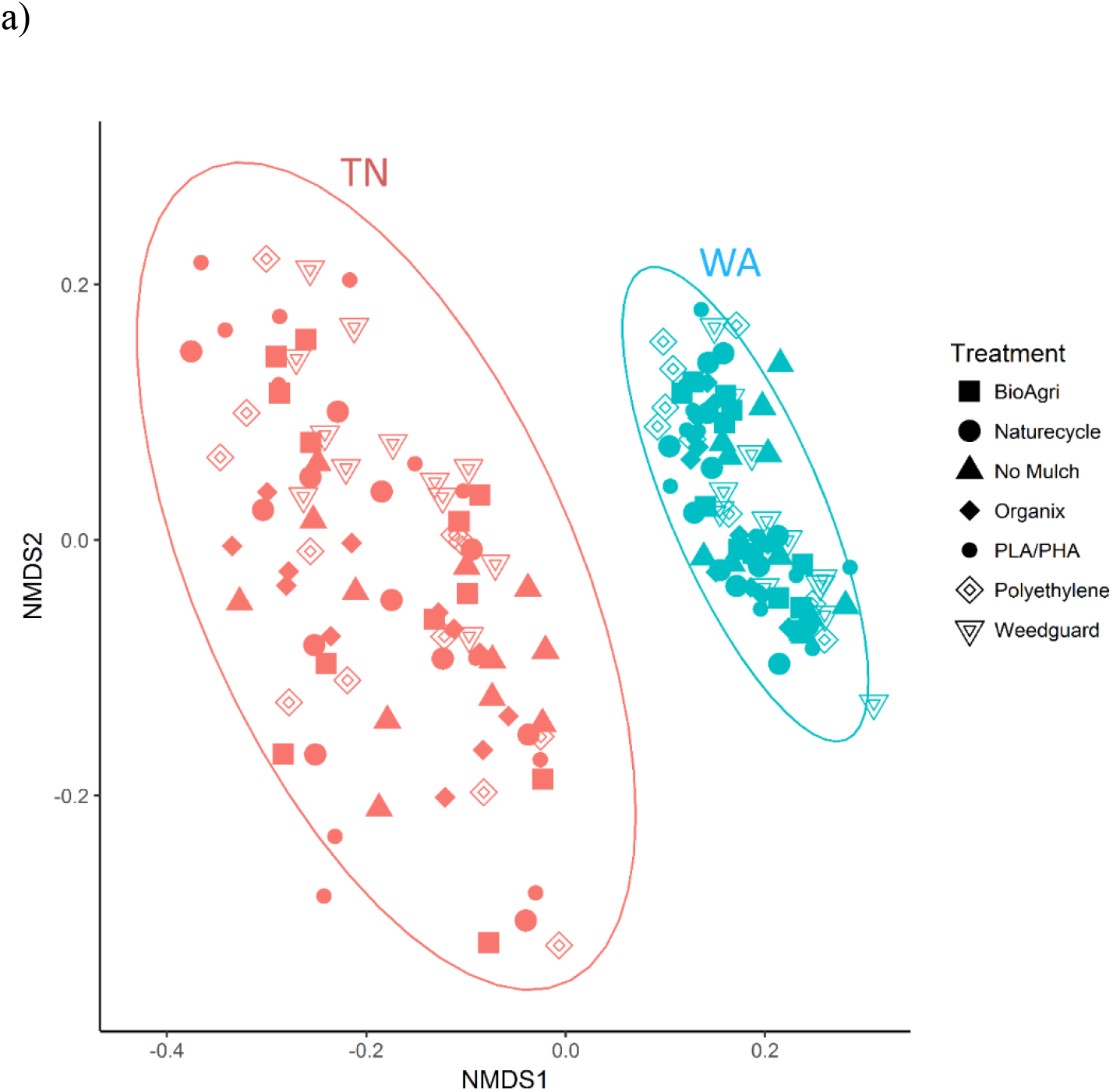
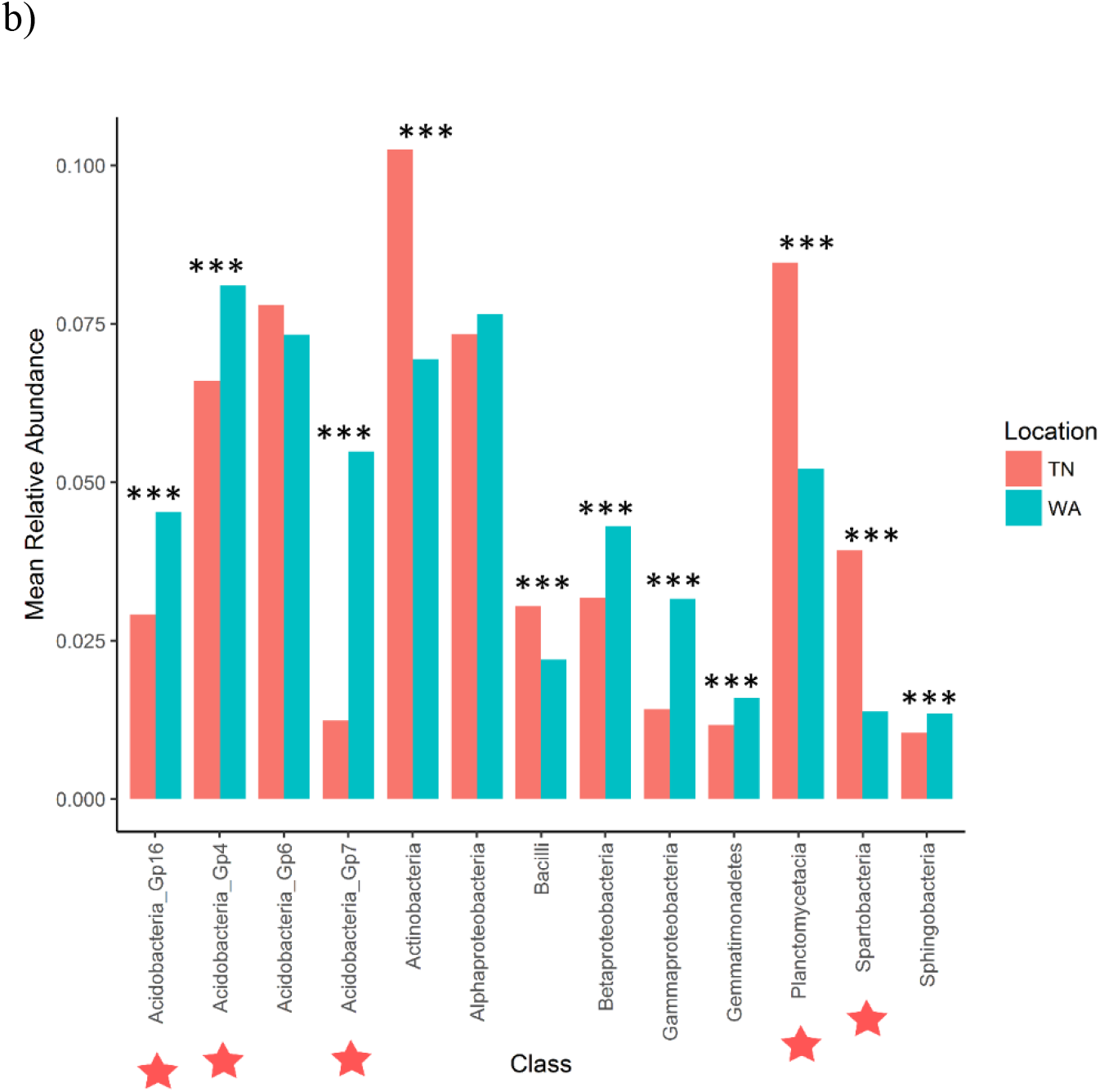
Bacterial community composition differences between the two field locations, showing communities from all four sampling times. a) Non-metric multidimensional scaling (NMDS) ordination of Bray-Curtis dissimilarities of OTU relative abundances, highlighting differences between location (PERMANOVA p = 0.001). Each point corresponds to the whole microbial community of one plot in the field (4 time points * 3 reps, total 12 points for each treatment). Ellipses denote clustering at 95% confidence. NMDS stress value: 0.14. b) Bar plot showing differences in mean relative abundance of the most abundant classes of bacteria in TN and WA, aggregating all treatments and all four sampling times. Asterisks denote significant differences between locations, determined by ANOVA (*p ≤ 0.05, **p ≤ 0.01, ***p ≤ 0.001). Red stars indicate taxa which cumulatively contributed up to 46% of the variance in microbial communities between TN and WA, determined using SIMPER.

**Table 3.**
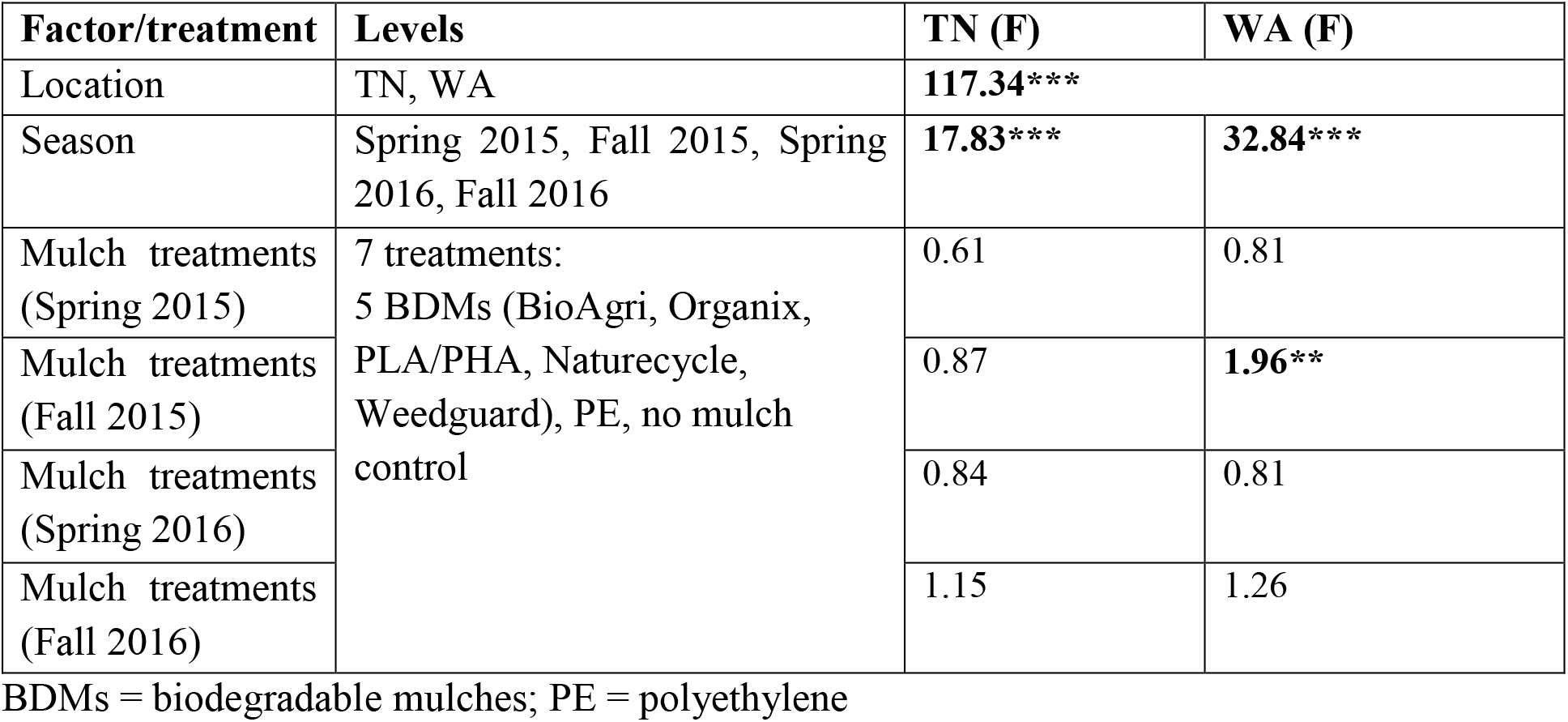
Results (F values) of PERMANOVA tests for differences in bacterial community composition by location (Knoxville (TN) and Mount Vernon (WA)), season and mulch treatment. Significant differences are in bold; *p < 0.05; **p < 0.01; ***p < 0.001

In addition to locational differences, bacterial communities also differed significantly between the different seasons (Table 3, Table S2). For both locations, Spring communities were more similar to each other than Fall communities (Fig 2a, b). SIMPER tests revealed that several genera of *Acidobacteria. Planctomycetaceae, Spartobacteria* and *Actinobacteria* (such as *Steptomyces sp.*) were cumulatively responsible for 60% of the seasonal variance in bacterial communities (Fig S2 and S3). Interestingly, *Streptomyces spp*. increased in percent relative abundance over time from Spring 2015 to Fall 2016 in both TN and WA (Fig S2).

**Fig 2.**
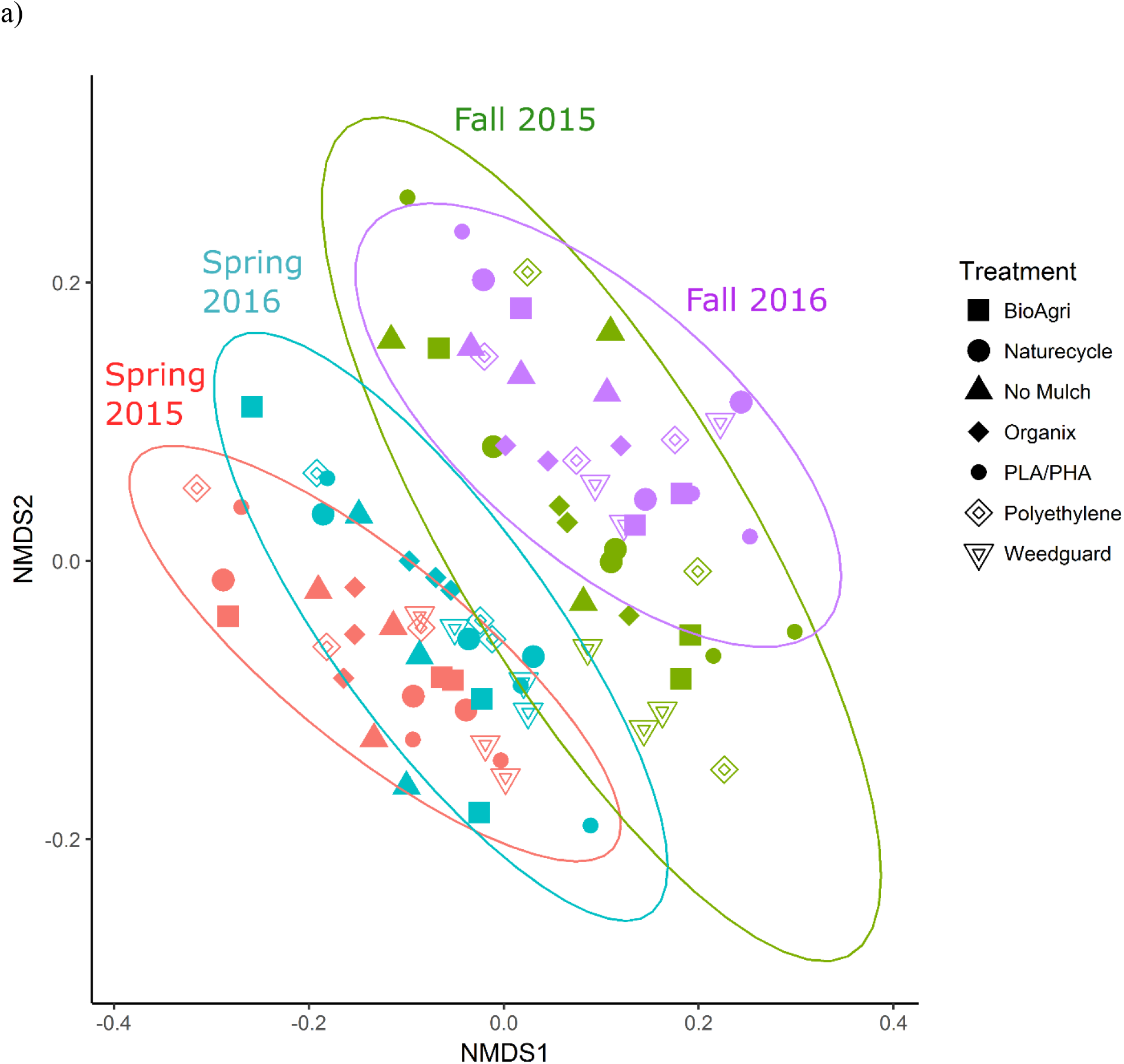
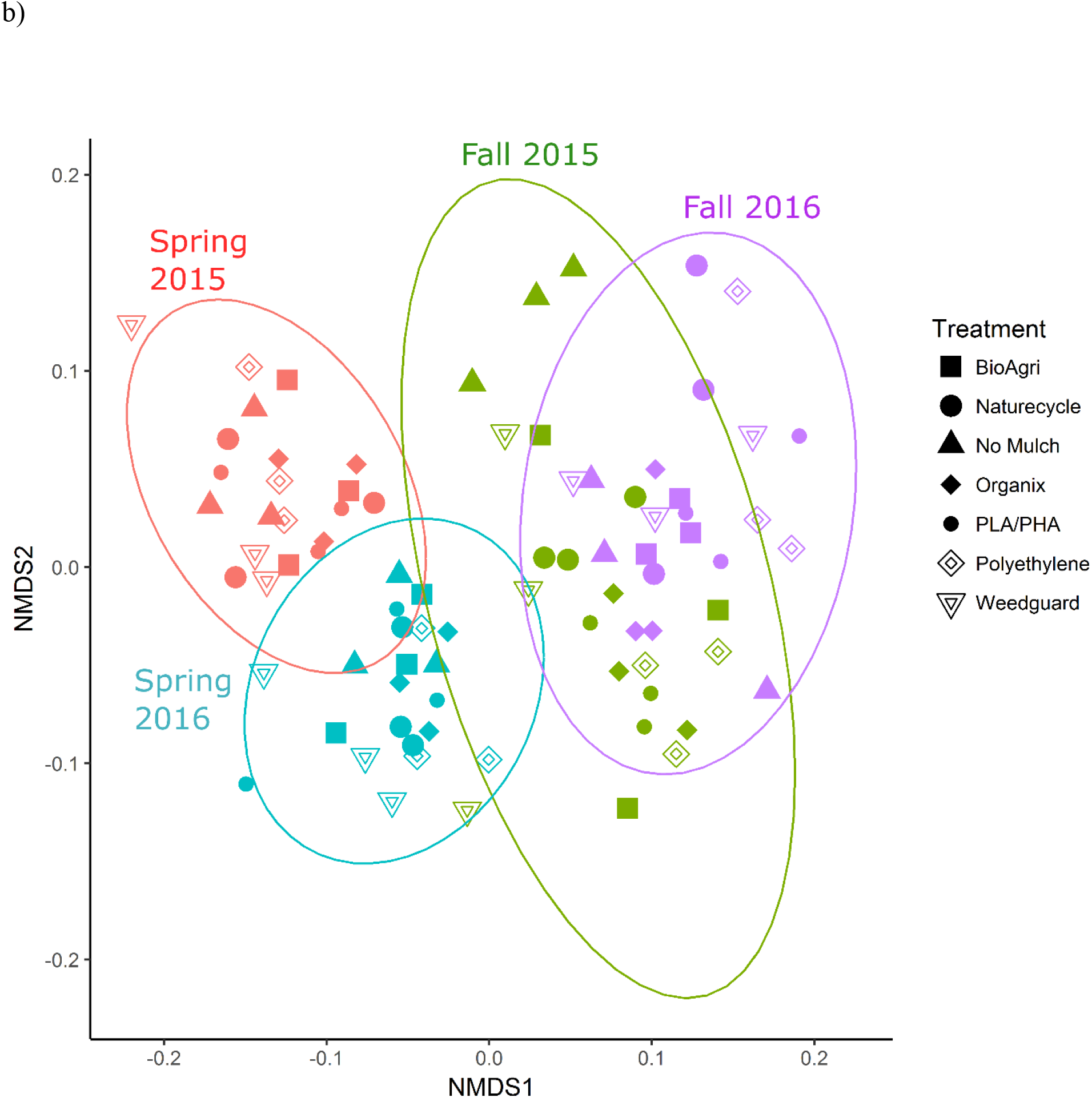
NMDS ordination of Bray Curtis dissimilarities of soil bacterial communities a) TN (p = 0.001) and b) WA (p = 0.001). Ellipses denote clustering at 95% confidence. NMDS stress value: 0.17 (TN), 0.16 (WA).

Unlike location and season, the mulch treatments did not have a significant effect on bacterial community structure (Fig 2). Because of the locational and seasonal differences, we additionally analyzed each time-location set separately, and did not detect any significant effects of treatment on community structure (Fig S4, Table 3, Table S2).

Alpha diversity of the soil bacterial communities was estimated using observed species richness and inverse Simpson index of diversity (Table S3). The observed species richness estimator measures count of unique OTUs in each sample. There were significant differences between TN and WA (p < 0.05) in richness estimates (Table 4, Fig 3a). TN had greater richness than WA throughout the experiment, ranging from 260 to 300 unique OTUs. WA richness estimates ranged from 250 to 280 OTUs over the two years. The locational differences in richness were due to a lower richness in Fall 2015, Spring 2016 and Fall 2016 in WA (Fig 3). The Inverse Simpson diversity index ranges were similar between TN and WA, ranging from 7 to 11.

**Fig 3.**
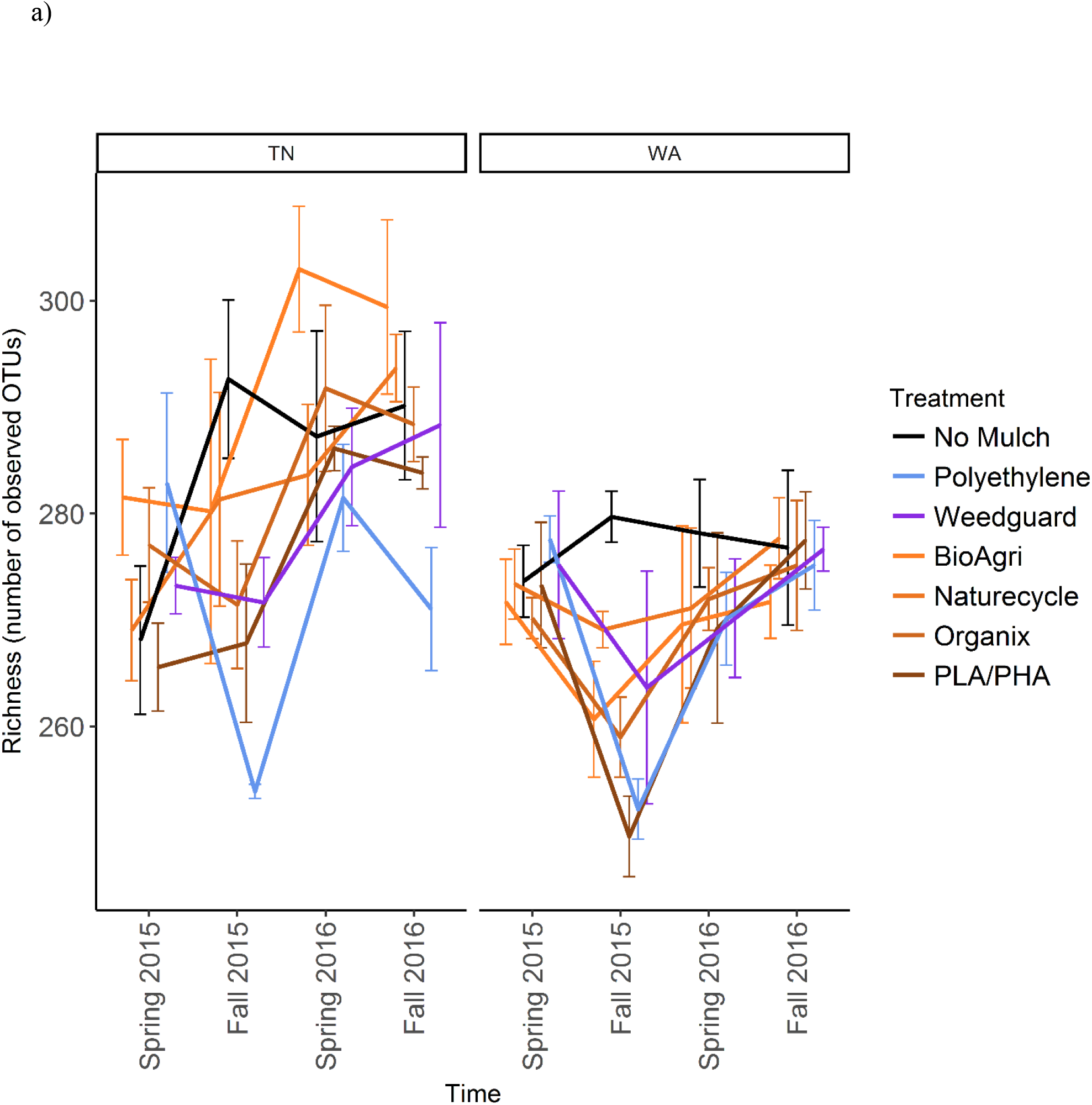
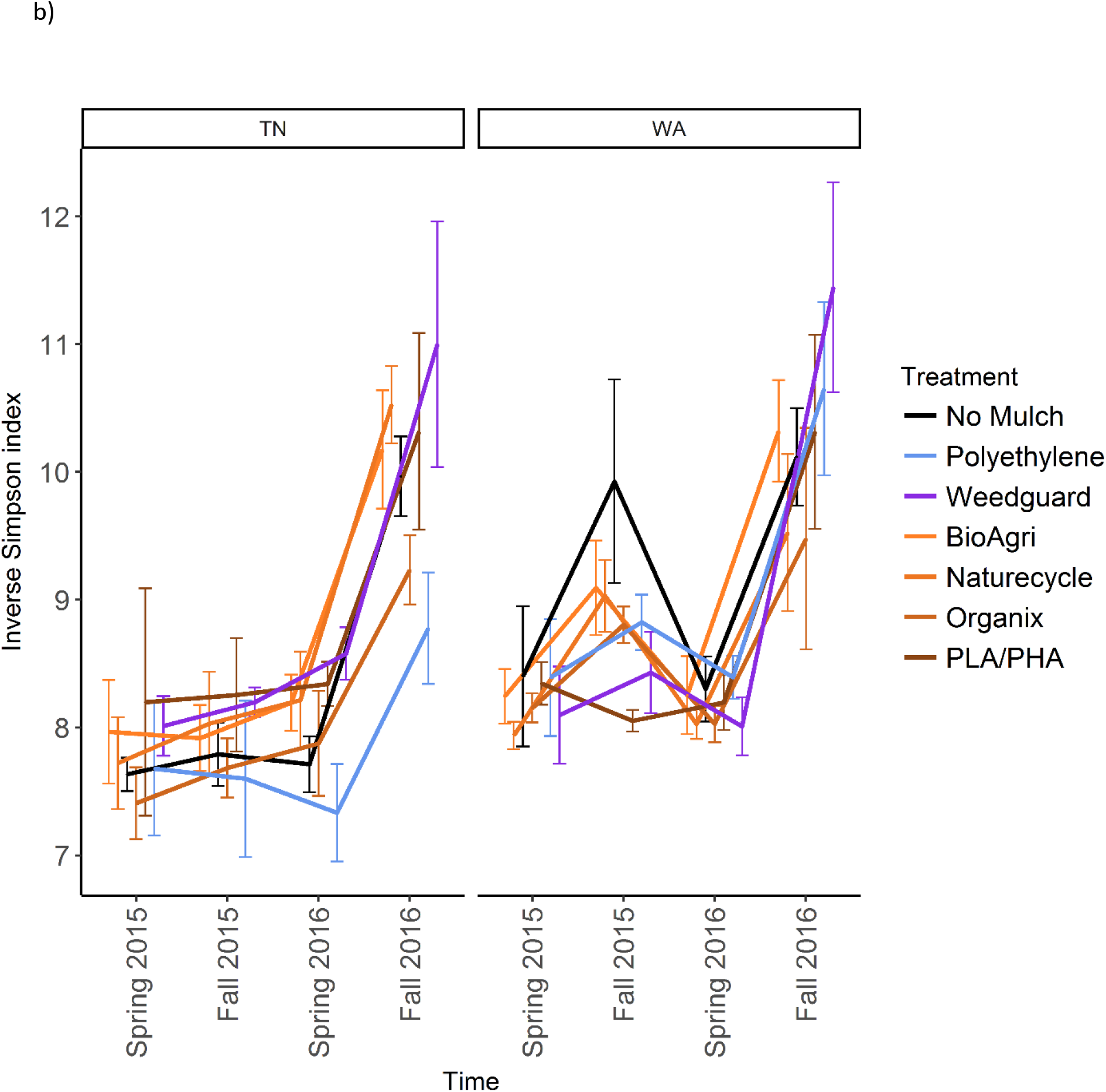
a) Richness (number of unique OTUs) and b) Inverse Simpson estimates over time of soil microbial communities in TN and WA. Error bars indicate SEM of three replicate samples.

**Table 4.**
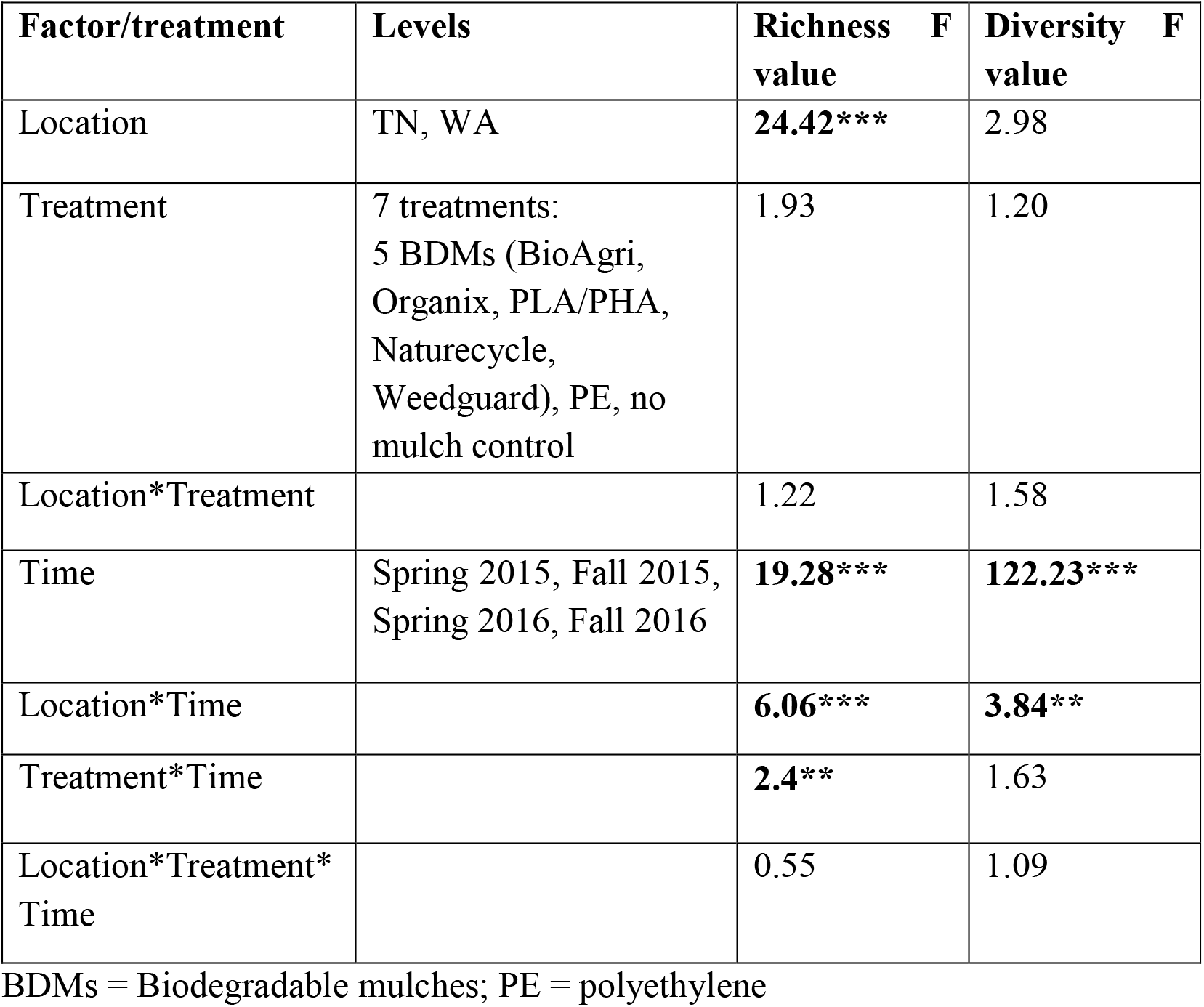
F values of fixed effects and interaction effects obtained from a mixed model analysis of variance of the alpha diversity metrics richness (number of observed OTUs) and diversity index (inverse Simpson) from Spring 2015 to Fall 2016 in Knoxville, TN and Mount Vernon, WA. Significant values are in bold, *p < 0.05; **p < 0.01; ***p < 0.001

For both TN and WA, there was a significant difference between seasons in both richness and inverse Simpson index (Table 4). The richness estimates in TN significantly differed between 2015 and 2016 (Fig 3a). In WA, Fall 2015 differed in richness from the other time points. In TN, Fall 2016 diversity was significantly higher than other seasons. Diversity estimates were significantly lower in Spring than in the Fall seasons for WA (Fig 3b).

In TN, PE had the lowest richness and BioAgri had the highest, however, treatment differences in richness estimates were not significant (Table 4) when analyzing data using a mixed model. Inverse Simpson diversity indices were also not significantly different between treatments (Table 4). Looking at the final time point in TN, diversity estimates were highest for Weedguard, and lowest for PE, and in WA, the estimates were highest for Weedguard, followed by PE with BDMs having lower diversity than PE or Weedguard, however these differences were not significant (Fig 3b).

### 3.3 Microbial community abundances

As a proxy for bacterial and fungal abundances, bacterial (16S) and fungal (ITS) rRNA gene copies were quantified using qPCR assays for soil samples from all seasons. In order to assess if gene abundances had significantly changed over the course of the experiment (Spring 2015 to Fall 2016) for each mulch treatment, a paired t-test was used to identify differences significantly different from zero (Table 5). There was a significant increase in bacterial gene copies under BDM and Weedguard treatments in WA, but no significant change for no mulch and PE treatments (Table 5). There was also a significant enrichment in fungal gene copies over time for two of the BDMs (PLA/PHA and Naturecycle) in WA. In TN, significant enrichment in bacterial gene copies was seen under Organix, PLA+PLA and PE treatments (Table 5) but no enrichment was seen in fungal gene copies. In order to determine if these changes were significantly different between treatments, the differences between the final (Fall 2016) and the initial (Spring 2015) abundances were analyzed using a mixed model analysis of variance in SAS v 9.3 and Tukey post hoc tests. In both locations, mulch treatments did not have a significant effect on the changes in either 16S or ITS gene copies over the course of the experiment (Fig 4a, b).

**Fig 4.**
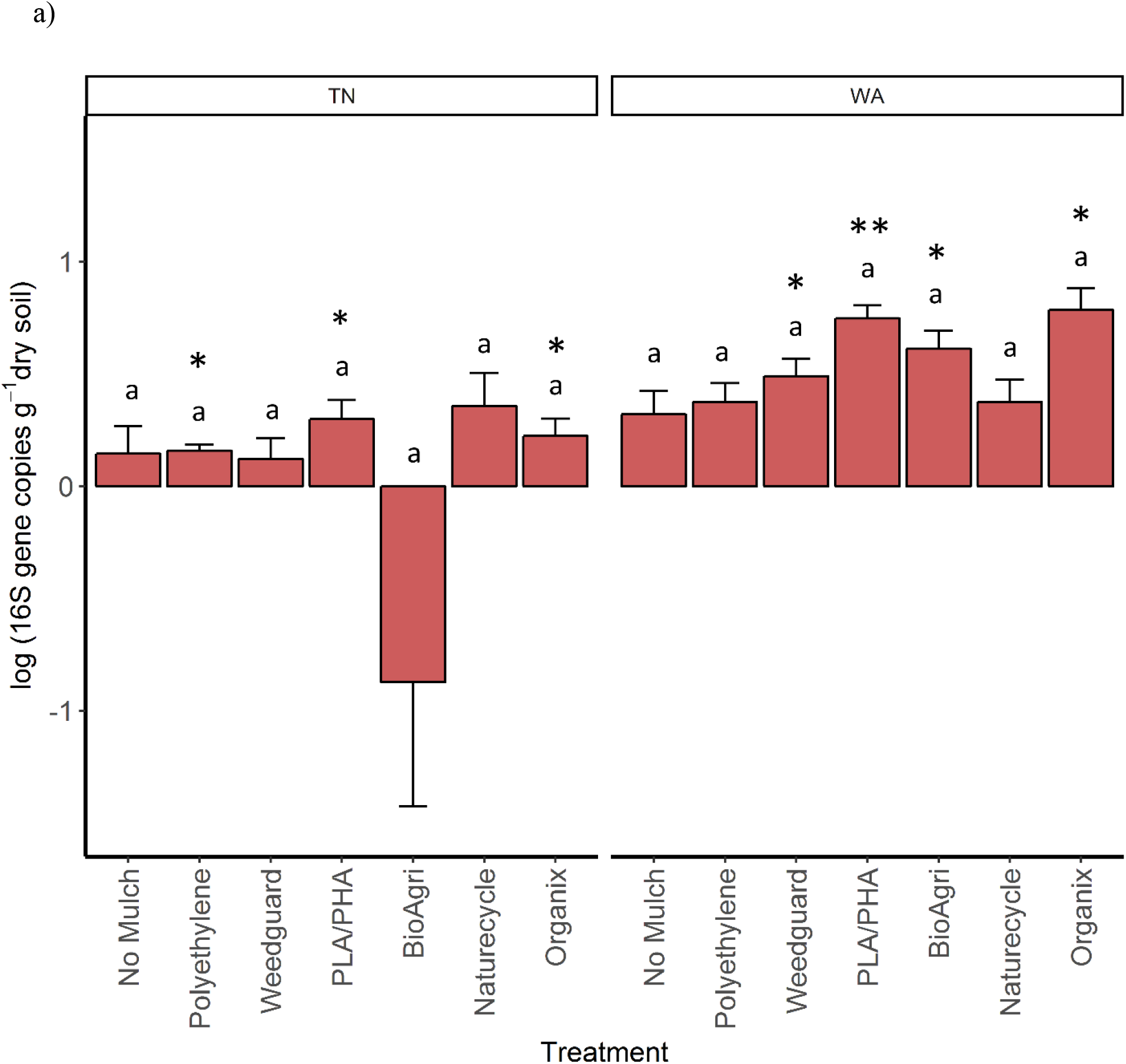
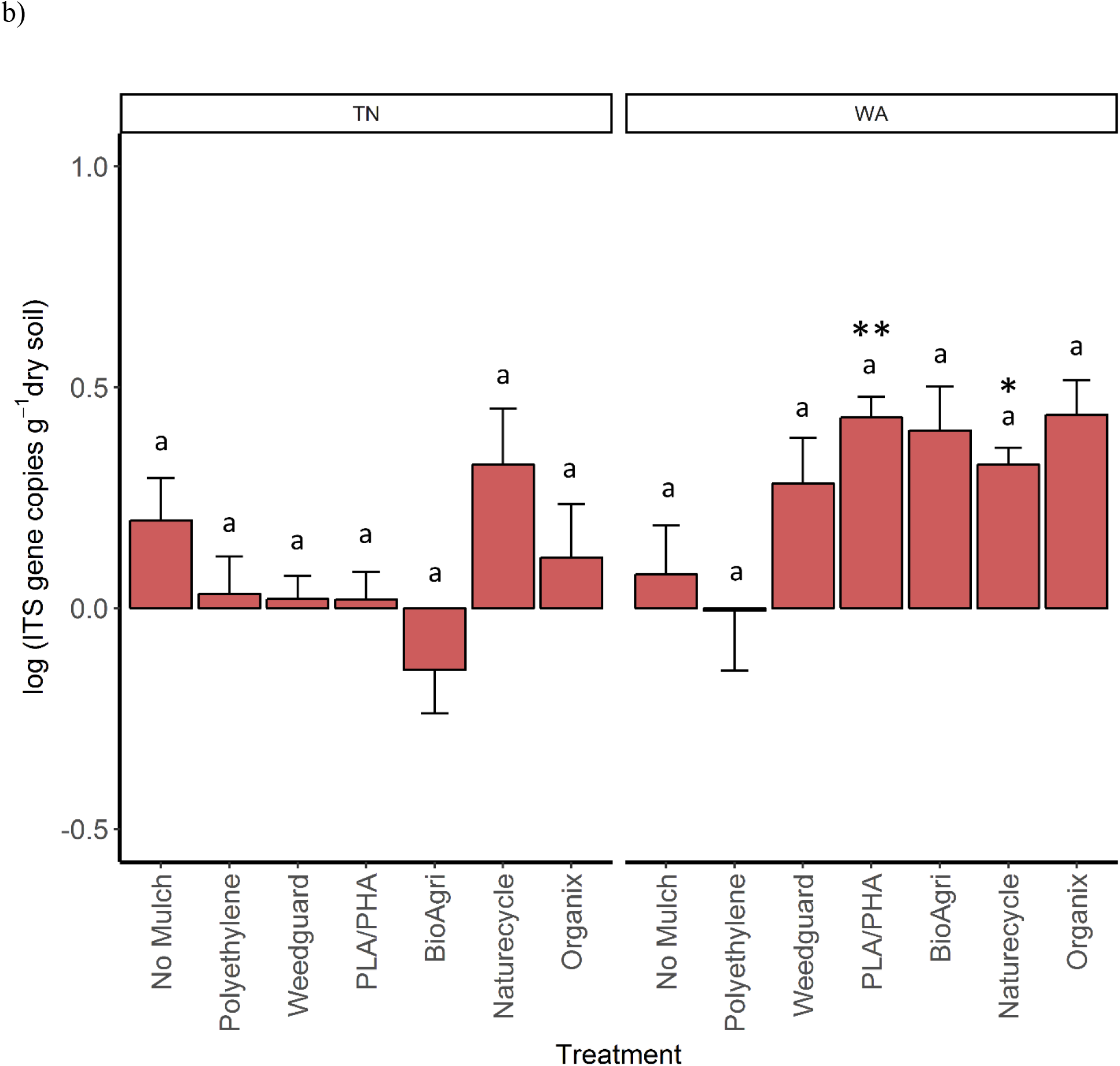
a) 16S and b) ITS gene copy number changes over two years in TN and WA analyzed using a mixed model analysis of variance. Final time point (Fall 2016) is plotted by subtracting baseline abundances from Spring 2015. Error bars indicate SEM of four replicate samples. Lowercase letters denote significant differences between treatments (p ≤ 0.05, Tukey’s HSD). Asterisks indicate treatments which showed significant enrichment using a paired t-test ((*p ≤ 0.05, **p ≤ 0.01, ***p ≤ 0.001).

**Table 5.** T values from paired t-tests comparing 16S and ITS initial abundances from Spring 2015 to final abundances from Fall 2016 to determine significant changes over the two-year experiment in Knoxville, TN and Mount Vernon, WA. Significant values are in bold, *p < 0.05; **p < 0.01; ***p < 0.001.

| Mulch Treatments and Controls |  | TN |  | WA |  |
| --- | --- | --- | --- | --- | --- |
|  |  | 16S | ITS | 16S | ITS |
| Treatments | BioAgri | 0.83 | 0.76 | **-4.30\*** | -2.27 |
|  | Naturecycle | -1.30 | -1.45 | -4.02 | **-6.87\*** |
|  | Organix | **-3.9\*** | -0.52 | **-4.30\*** | -3.05 |
|  | Experimental PLA/PHA | **-3.51\*** | -0.20 | **-8.34\*\*** | **-5.64\*\*** |
|  | Weedguard | -0.80 | -0.89 | **-3.52\*** | -1.50 |
| Controls | Polyethylene | **-4.06\*** | -0.23 | -2.53 | -0.02 |
|  | No mulch | -0.65 | -1.21 | -1.74 | -0.38 |

### 3.4 Microbial community functions

To assess potential functional responses of the soil microbial communities, extracellular enzyme potential rate assays were conducted for common carbon, nitrogen, and phosphorus cycling enzymes in soil (Table 2). The data were combined over the two years to visualize Bray Curtis similarities of the enzyme rate profiles (Fig 5). Locational differences in the enzyme profile were significant (p < 0.05), as were seasonal differences in both TN (p < 0.05) and WA (p < 0.05) evaluated using PERMANOVA (Fig 5). However, mulch treatment did not have a significant effect on the enzyme profile for any of the seasons at either location (p < 0.05). NMDS ordination for the final sampling time point Spring 2017 is shown in Fig S5, showing no clear treatment differences.

**Fig 5.**
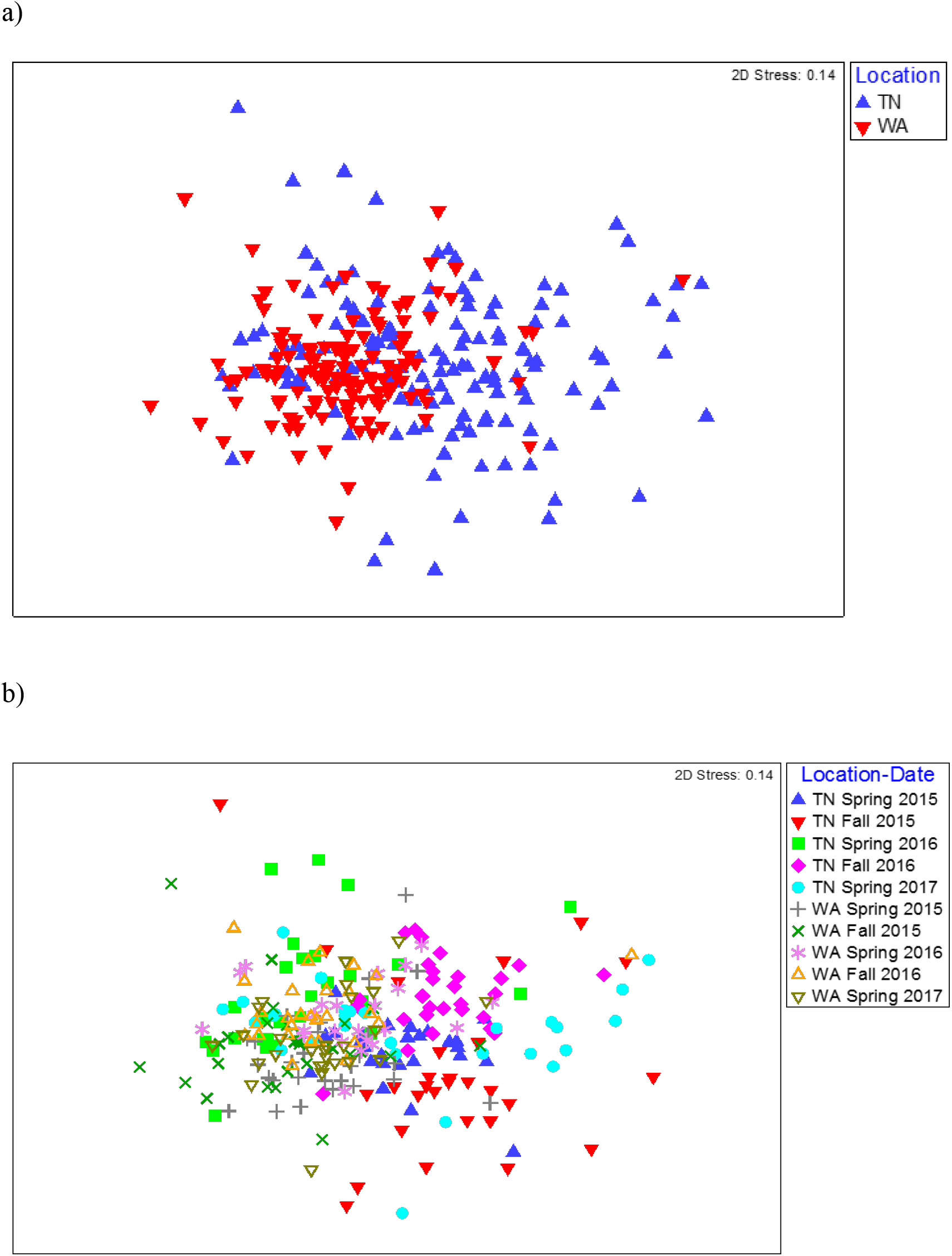
NMDS ordination depicting Bray-Curtis similarity of the functional profile of soil microbial communities (based on 7 soil enzyme activity rates) across a) Location (p < 0.05) and b) Date (p < 0.05).

In general, the enzyme activity rates oscillated between higher activities in the Spring and lower activities in the Fall. When analyzed separately for each enzyme, the data over the two years revealed a significant effect of sampling time in TN for all seven enzymes assayed. In WA, enzyme activities of β-xylosidase, β-glucosidase, α- glucosidase, N-acetyl β glucosaminidase and phosphatase were significantly different between sampling times (Fig 6). In WA, cellobiosidase and leucine amino peptidase activities remained unchanged across the seasons (10-22 nmol activity g^-1^ dry soil h^-1^ for cellobiosidase and 200-375 nmol activity g^-1^ dry soil h^-1^ for leucine amino peptidase) (Fig 6).

**Fig 6.**
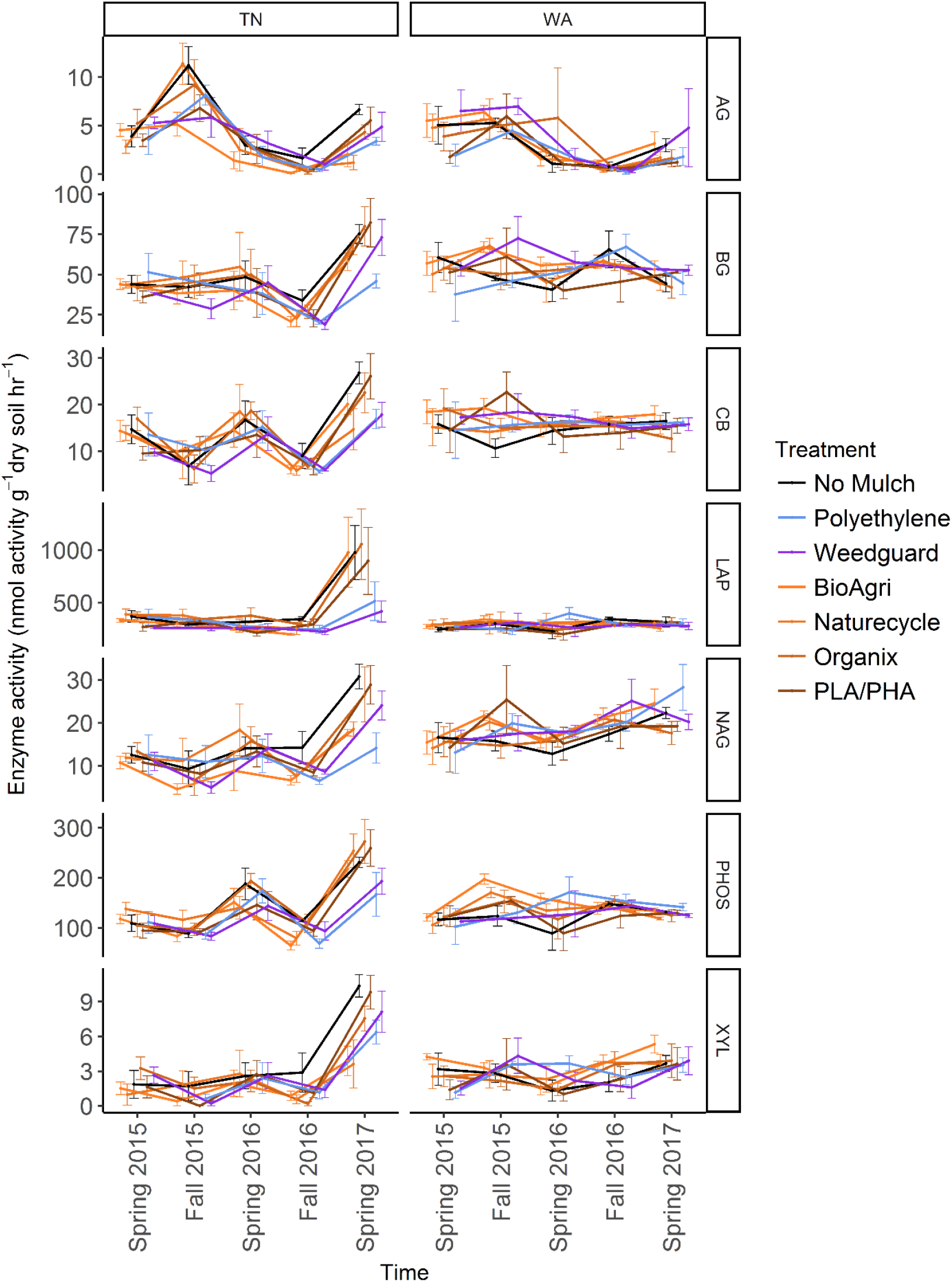
Changes in soil enzyme activity over time across mulch treatment (p values reported in Table 6). Error bars indicate SEM of four replicate samples.

**Table 6.** F values of fixed effects and interaction effects obtained from a mixed model analysis of variance of the soil enzyme activities from Spring 2015 to Spring 2017 in Knoxville, TN and Mount Vernon, WA. Significant values are in bold, *p < 0.05; **p < 0.01; ***p < 0.001

|  | TN |  |  | WA |  |  |
| --- | --- | --- | --- | --- | --- | --- |
|  | Treatment*Time<br>(F) | Time (F) | Treatment (F) | Treatment*Time<br>(F) | Time (F) | Treatment (F) |
| Effect |  |  |  |  |  |  |
| β-xylosidase | 1.55 | **46.48\*\*\*** | 2.21 | 0.88 | **5.12\*\*\*** | 0.89 |
| β-glucosidase | 0.92 | **29.56\*\*\*** | 1.49 | 0.91 | **3.44\*** | 0.84 |
| α-glucosidase | 1.52 | **40.16\*\*\*** | 2.62 | 0.65 | **13.31\*\*\*** | 1.12 |
| N-acetyl $\beta$<br>glucosami-<br>nidase | 1.26 | **34.60\*\*\*** | **2.53\*** | 0.78 | **6.06\*\*\*** | 0.64 |
| β-D<br>cellubiosidase | 0.88 | **32.82\*\*\*** | 1.03 | 0.72 | 0.27 | 0.75 |
| Phosphatase | 1.04 | **68.23\*\*\*** | 1.37 | 0.96 | **4.10\*\*** | 1.13 |
| Leucine amino<br>peptidase | 0.96 | **28.83\*\*\*** | 1.71 | 0.77 | 0.65 | 0.34 |

When averaged across seasons, mulch treatment differences were not significant for any soil enzymes in WA (Table 6). However, in TN, an effect of mulch treatment was observed for N-acetyl β glucosaminidase activities (Table 6). N-acetyl β glucosaminidase activity was reduced under BDMs and PE compared to no mulch plots. Interaction effects of mulch treatment and time of sampling were not detectable for any of the enzymes assayed in TN or WA (Table 6).

## 4. Discussion

Characterizing the soil microbial communities under the different biodegradable mulches and non-biodegradable PE mulch revealed no significant effect of mulch type on bacterial community structure. This is in contrast to other studies that have reported altered bacterial communities in soils under BDMs (29, 40, 41), and under non-biodegradable plastic mulches (42, 43). Such opposite findings could be due to differences in methodology: for example, the studies by Koitabashi et al. (40) and Muroi et al. (41) were shorter laboratory incubation studies in controlled conditions (28 to 30°C), used pure polymer feedstock (rather than commercial film formulations which include plasticizers and other additives) and relied on detection methods such as polymerase chain reaction-denaturing gradient gel electrophoresis (PCR-DGGE). Laboratory studies under controlled conditions often result in more rapid microbial responses to treatments as opposed to field studies where variable environments introduces more noise. Our lack of observed difference may also be because of a realistic, but low, plastic to soil ratio: for example, Muroi et al. (41) used soil burial studies with an artificially high 1.8 g PBAT films in 300 g soil with added 30 ml of basal medium. Finally, our aim was to characterize responses in bulk soil communities to understand the overall system level response to plastic films, so we likely missed changes happening on smaller spatial scales. For example, Li et al. (29) reported changes in microbial communities in soils that were sampled in close proximity to buried mulch films, indicating that microbial communities in the immediate vicinity of the films may be affected. Here we show that any local effects of mulch films are not detectable at a plot/field scale, at least over a 2-year period.

We did note significant differences in soil bacterial composition by location and season, which has been observed in other studies (29, 30), confirming that mulch effects are minimal compared to other drivers of community structure variation. It is well accepted that local soil conditions such as temperature, moisture and pH play a pivotal role in shaping microbial communities (30, 44, 45).

In this study, the location differences in communities were attributed to higher relative abundances of *Acidobacteria, Actinobacteria* and *Planctomycetes* in TN and higher abundances of *β-* and *γ- Proteobacteria* in WA. This corresponds with higher pH and saturated K in TN and higher soil organic matter and soil moisture in WA. Both pH and water content are major edaphic factors that influence temporal and spatial variation in soil microbial communities (44, 46). Changes in soil physicochemical properties and different climates and soil types between TN and WA could explain such locational differences. Seasonal differences in communities were driven by significantly increased percent relative abundance of *Acidobacte*r *Gp6, Gp4* and *Gp7* in Spring in TN as compared to Fall. Additionally, significantly greater abundances of *Planctomycetaceae* and *Streptomyces* were seen in Fall compared to Spring in TN. In WA, *Acidobacteria_Gp6* and *Spartobacteria* showed significantly greater percent abundances in Spring compared to Fall whereas *Streptomyces* sp. showed significantly higher percent abundance in Fall compared to Spring (Fig S2). Seasonal tillage operations often reset many of the soil properties which can explain why the abundances of some taxa oscillated between Spring and Fall. *Streptomyces sp.* belongs to class Actinobacteria and have demonstrated polymer degrading capabilities (47). However, because we did not observe differences in the relative abundance of this taxa between BDMs, PE or no mulch control, this increase is likely attributable to the agronomic management of the plots (e.g. plant species, irrigation or fertilizer regimes etc.).

Richness estimates showed significant differences across locations and seasons in TN and WA. Diversity estimates were only significantly different between seasons, but not location. Mulch materials did not have a consistent impact on bacterial richness or diversity. A previous study evaluating microbial diversity using PCR-DGGE showed no difference in ammonia oxidizer diversity under biodegradable and non-biodegradable mulching materials one year after tilling plastics into soil (48). The higher richness estimates under BDMs compared to PE treatments, which was significant in Fall 2015 in WA, suggested that tilled BDMs may help promote richness in the soil environment. Increased warming potential under PE mulch could also contribute to suppression of microbial activity which may also have impacts on community richness or diversity.

Gene copy abundances in soils were used as a proxy for bacterial and fungal abundances. In WA we observed an enrichment of both bacteria and fungi under BDM and Weedguard treatments over the course of the two-year experiment. In TN, we observed bacterial, but no fungal, enrichment in two of the four BDM plots and PE plot. A mixed model analysis of changes over the course of the experiment was not able to detect significant differences between treatments. It should be noted that enrichment was observed under BDM but not the PE plots in WA, suggesting that this is response to the incorporation of BDMs into the soil (as opposed to an indirect effect of microclimate modification, such as soil warming). Previous studies have also demonstrated increased fungal abundances in soil because of BDM incorporation (29, 41, 49, 50). Fungi have been observed to be important colonizers and degraders of BDMs (30, 40, 41). Tilled into soil, BDMs are a very small input of carbon when taking into account the volume of soil into which they are incorporated (5). For comparison, the input of mulch carbon added to the soil in this study was a significantly smaller amount (6-25 g C m^-2^) (51) compared to the amount added from cover crop residues (142 g C m^-2^) (52). However, the growth of soil microbes in agricultural soil is usually carbon-limited and several studies have demonstrated responses by soil microbes to these small inputs (5).

There is also precedent for the differential responses in microbial enrichment we observed between the two locations, with both fungal and bacterial enrichment in WA, but only bacterial enrichment in TN. In a similar study comparing BDM effects in three locations, it was found that BDMs resulted in soil fungal enrichment in Texas and bacterial enrichment in TN (29). In one study, soil pH was shown to be the best predictor of bacterial community composition across different land use types, while fungal communities were shown to be most closely associated with changes in soil nutrient status such as extractable P concentrations and C:N ratios (53). Both TN and WA soils had comparable fungal gene abundances initially in the Spring of 2015. However, since the microbial communities in WA were seen to be controlled by the presence of organic matter (Fig S1) and WA soils had higher C:N ratios than TN soils this could have contributed to a fungal enrichment in WA but not in TN.

Enzyme assays were conducted to assess potential activity rates for common carbon, nitrogen and phosphorus cycling enzymes in soil. As with bacterial community structure, enzyme activity profiles showed the greatest differences by location and season (Fig 5, Table 6). The seasonal oscillation in enzyme activities seen for almost all the enzymes could be attributed to seasonal tillage operations which tend to offset many of the soil biological functions (54–56) (Fig 6). This was also observed for many of the soil physicochemical properties (31). Mulch treatments had significant effects on N-acetyl-β -glucosaminidase (NAG) in TN. NAG was decreased under mulches compared to no mulch treatments, with the greatest decrease observed under PE. NAG catalyzes the hydrolysis of chitin to form amino sugars which are major sources of mineralizable nitrogen in soils and thus is important in carbon and nitrogen cycling in soils. Xylosidase activity was also reduced under mulch treatments compared to no mulch plots in TN though not significant. Because we saw decreases under all mulch treatments for NAG in TN, this is likely an indirect effect of the mulches via microclimate modification, rather than a direct effect of mulch fragments tilled into the soil. All mulches warm the soil, with PE often having a greater soil warming potential compared to BDMs (2, 57). Mulches also increase soil moisture levels (58). Consequently, changes in soil temperature and moisture will affect enzyme pool sizes (59). The reduction in activity under plastic mulches may be because TN has a warmer climate where plastic mulches can push temperatures above optima limiting soil microbial activity (57). Mean soil temperatures in summer under mulched plots were 24.7 °C at 10 cm depth in TN, whereas in WA it was 18.7 °C. Un-mulched plots had mean summer soil temperatures of 23.8 °C for TN and 17.0 °C for WA (31). In the month of June in both years, soil temperatures exceeded 30 °C under mulched plots in TN, but were less than 30 °C for no mulch plots. It has been reported that fungal and bacterial growth rates have optimum temperatures around 25 to 30 °C in agricultural and forest humus soils, while at higher temperatures lower growth rates are found (60). This decrease in growth rate was shown to be more drastic for fungi than for bacteria, resulting in an increase in the ratio of bacterial to fungal growth rate at higher temperatures. Thus, the high temperatures under mulches in the summer in TN were above optimum growth conditions for soil microbes and may have reduced soil enzyme activities. Cold-adapted microorganisms, which are expected to be more prevalent at the WA site, tend to respond more efficiently to increased temperature than warm-adapted microbes (61). The greatest relative temperature sensitivity of decomposition processes has been observed at low temperatures (62). Warming experiments have revealed reduced xylosidase activity in soils (5-15 cm deep) under medium-warmed plots compared to unwarmed plots (59). It has also been reported that warming induces decreases in the temperature sensitivity of β-xylosidase activity in the H horizon (63). One study reported greater increase of the relative temperature sensitivity of XYL and NAG (important for C cycling) at lower temperatures, compared to amino peptidase enzymes suggesting that temperature plays a pivotal role in regulating the use of substrates. Thus, the turnover of easily degradable C substrates (like glucose) is more sensitive to temperature than higher molecular compounds, at least for cold soils (64).

Looking specifically at studies which assessed soil enzyme activities after treatment with biodegradable plastic film, one field study reported that soil microbial biomass and beta-glucosidase activity were most responsive to mulch; however that study did not have PE as a control, so it is unclear if this response was specific to BDMs or just related to plastic mulching generally (65). That study also focused on soils in close proximity to plastic, rather than bulk soil responses. Laboratory studies have shown increased esterase activity in soils during the degradation of PBSA (66), and increased microbial activity as per a fluorescein diacetate hydrolysis test during the degradation of a variety of biodegradable polymers (67). These studies provide insight into the potential of these enzymes in the degradation process of BDMs. Other studies that have looked at more general activity responses by microbes under plastic mulches (i.e. respiration) have reported mixed results: some have observed increases in activity under plastic mulches (68–71), while others report decreased activities (57).

In our recent paper from the same field sites as mentioned in the present study we have shown that biodegradable mulches do not have a significant impact soil health in terms of a suite of soil quality parameters tested over two years in TN and WA (31). Our findings corroborate the results from Sintim et al. (31) where it has also been shown that locational and seasonal variations are more important drivers of change in overall soil health under BDM tillage operations as compared to mulch treatment itself.

## 5. Conclusion

Two years of biodegradable and PE mulch treatments in a vegetable agroecosystem in two locations revealed some minor effects on soil microbial communities and their functions. While we were not able to detect any significant effect of plastic mulches on bacterial community structure, richness or diversity, we did observe other impacts on the communities that were location-dependent. In particular, we noted that in WA, biodegradable mulches enriched for both bacteria and fungi, suggesting a response to BDM incorporation into soils; in contrast only bacterial enrichment was apparent in TN, and only for three of the five plastics tested. We additionally observed decreases in specific enzyme activities (NAG) under mulch treatments in TN but not WA, which may be attributable to increased temperatures under the plastics (i.e. microclimate modification) rather than mulch fragment incorporation into soil. Together, this shows that plastic mulches do have minor impacts on soil microbial communities and their functions, and that BDMs may have effects different from PE plastic mulches. As microbes are the drivers of soil carbon and nutrient cycling, changes in bacterial and fungal abundances and/or activity can have repercussions for soil organic matter dynamics and nutrient availabilities. Longer term studies of repeated BDM incorporation are needed to determine if these microbial responses will significantly affect soil functioning and health. In addition, the fact that we saw different responses by the communities in two locations under identical management may mean that the ultimate impact of plastic mulching on soil functioning may be dependent on local climate and soil conditions.

## 6. Acknowledgements

This work was supported by the United States Department of Agriculture Specialty Crops Research Initiative, CAP (Award 2014-51181-22382 to JMD). Field experiments were designed and managed by A. Wszelaki, C. Miles, D. Ingles and D. Hayes, with help from staff at the East Tennessee Research and Education Center (Knoxville, TN) and Northwestern Washington Research and Extension Center (Mount Vernon, WA). We are grateful to BioBag Americas, Inc. (Palm Harbor, FL, USA), Organix Solutions (Maple Grove, MN, USA), Custom Bioplastics (Burlington, WA, USA), (Metabolix Inc., (Cambridge, MA, USA), and Sunshine Paper Co. (Aurora, CO, USA) for the donation of mulches for the research experiments, and Techmer PM (Clinton, TN, USA) for preparation of the carbon black dye masterbatch and compounding of the PLA/PHA formulation used to prepare the PLA/PHA mulch film. Arnold Saxton provided statistical advice. We thank M English, J Moore, S Schexnayder, M Valendia, S Schaeffer, M Anunciado, K Henderson, S Keenan, LS Taylor, J Liquet, Shuresh Ghimire, Andy Bary, M Starrett, D Cowan-Banker and other members of the JMD and MF labs for help with soil sample collection and processing. M Flury and D Hayes provided critical feedback on the manuscript.

